# Simple and practical sialoglycan encoding system reveals vast diversity in nature and identifies a universal sialoglycan-recognizing probe derived from AB_5_ toxin B subunits

**DOI:** 10.1101/2021.05.28.446191

**Authors:** Aniruddha Sasmal, Naazneen Khan, Zahra Khedri, Benjamin P. Kellman, Saurabh Srivastava, Andrea Verhagen, Hai Yu, Anders Bech Bruntse, Sandra Diaz, Nissi Varki, Travis Beddoe, Adrienne W. Paton, James C. Paton, Xi Chen, Nathan E. Lewis, Ajit Varki

**Author notes:** Address correspondence to: Ajit Varki, UCSD School of Medicine, La Jolla, CA 92093-0687.

## Abstract

Vertebrate sialic acids (Sias) display much diversity in modifications, linkages and underlying glycans. Slide microarrays allow high-throughput explorations of sialoglycan-protein interactions. A microarray presenting ∼150 structurally-defined sialyltrisaccharides with various Sias linkages and modifications still poses challenges in planning, data sorting, visualization and analysis. To address these issues, we devised a simple 9-digit code for sialyltrisaccharides with terminal Sias and underlying two monosaccharides assigned from the non-reducing end, with three digits assigning a monosaccharide, its modifications, and linkage. Calculations based on the encoding system reveals >113,000 likely linear sialyltrisaccharides in nature. Notably a biantennary *N*-glycan with two terminal sialyltrisaccharides could thus have >10^10^ potential combinations and a triantennary *N*-glycan with three terminal sequences, >10^15^ potential combinations. While all possibilities likely do not exist in nature, sialoglycans encode enormous diversity. While glycomic approaches are used to probe such diverse sialomes, naturally-occurring bacterial AB_5_ toxin B subunits are simpler tools to track the dynamic sialome in biological systems. Sialoglycan microarray was utilized to compare sialoglycan-recognizing bacterial toxin B subunits. Unlike the poor correlation between B subunits and species phylogeny, there is stronger correlation with Sia-epitope preferences. Further supporting this pattern, we report a B subunit (YenB) from *Yersinia enterocolitica* (broad host range) recognizing almost all sialoglycans in the microarray, including 4-*O*-acetylated-Sias not recognized by a *Y. pestis* orthologue (YpeB). Differential Sia-binding patterns were also observed with phylogenetically-related B subunits from *Escherichia coli* (SubB), *Salmonella* Typhi (PltB), *S*. Typhimurium (ArtB), extra-intestinal *E*.*coli* (*Ec*PltB), *Vibrio cholera* (CtxB), and cholera family homologue of *E. coli* (EcxB).

## Introduction

Glycans play critical roles in numerous structural and modulatory functions in cells. Cell surface glycans are particularly important as they are involved in various biological processes, like cell signaling, host-pathogen interaction, and immune interactions (1, 2). Sialic acids (Sias) are nine-carbon monosaccharides that are the most prominent terminal monosaccharides found in mammalian cell surface glycomes. The sialome is defined as the ‘‘total complement of sialic acid types and linkages and their mode of presentation on a particular organelle, cell, tissue, organ or organism—as found at a particular time and under specific conditions’’(3). The huge diversity in sialomes arises from the monosaccharide identity, functional modification, and various linkages of Sias. Besides the two most common mammalian Sias, *N*-acetylneuraminic acid (Neu5Ac) and *N*-glycoylneuraminic acid (Neu5Gc), the Sia family includes ketodeoxynononic acid (Kdn), and neuraminic acid (Neu), which are rarely found in mammalian glycomes.

Glycans are one of the major classes of macromolecules in nature. Due to the structural complexity of glycans compared to the simpler linear sequences of nucleotides and peptides, the development and availability of glycan informatics resources, such as user-friendly nomenclature, structure motif finder, and database are still limited, hindering the advancement of glycan research (4, 5). Unlike nucleic acids and proteins, glycan sequences are also comprised of a large number of possible monosaccharide residues, variable linkages, and branching possibilities (6, 7). The possibility of multiple functional modification and linkages on seven out of the nine carbons in Sias made it particularly challenging to establish a functional yet convenient representation of sialosides. The diversity of sialosides increases further when the underlying glycan structures are taken into account.

To partly address this issue, an abbreviated text-based nomenclature describing the glycan structure was established by the IUPAC-IUBMB (International Union for Pure and Applied Chemistry and International Union for Biochemistry and Molecular Biology) (8). However, this three-letter coding system is ambiguous and not easily transferable to a machine-readable format. Later LINUCS (Linear Notation for Unique description of Carbohydrate Sequences) was developed to address the IUPAC limitations (9), but it has not had universal acceptance and applications. Several efforts have since been made to develop a comprehensive representation of complex glycan structures (10–13) and their associated reaction rules (14), and establish an algorithm for the mining of high-complexity glycan motifs involved in biological processes (15). Early attempts have been made to implement useful identifiers for explicitly describing the glycan structures in databases like Complex Carbohydrate Structure Database, CarbBank, CFG (http://www.functionalglycomics.org/glycomics/publicdata/primaryscreen.jsp.), CarbArrayART (https://glycosciences.med.ic.ac.uk/carbarrayart.html) etc. (14, 16–20). Recently, assignment of unique glycan structure identifiers in the GlyTouCan repository (21) has been increasingly accepted by the glycomics community. A graphical representation of glycan structures by the ‘Symbol Nomenclature for Glycans’ (SNFG) system has also been standardized (22).

Automated and high-throughput analytical techniques, such as glycan microarray produces a large amount of glycan binding data (23, 24). Considerable efforts have been made to establish proper nomenclature and motif finders for in-depth study of structure and function relationships of glycans (25–27). Many recent studies have focused in glycan motif discovery to bring generalizable meaning to lectin array analysis (15, 26, 28), glycan database analysis (29–31) and glycoprofile analysis (32). Functional meaning or biosynthetic feasibility discovered in the existence or enrichment of one motif can be used to create generalizable expectations regarding the functional importance or biosynthetic feasibility of a novel glycan (33). Historically, our groups have been focusing on identifying novel biological roles of terminal Sias and consequently, and over the years we developed a sialoglycan-focused microarray platform to explore the novel biomolecular interactions with the sialosides (34–36). Our extensive sialoglycan inventory includes mostly sialyltrisaccharides with various modification and linkages of Sia. The ever-increasing sialoglycan microarray library demands an efficient coding-based structure representation system to streamline the printing, sorting, analysis, and motif-finding processes. The available tools for the systematic analysis of glycan microarray data do not address the variability of the terminal sialic acids on the mammalian glycans (37). A coding-based systematic representation of complex *N*-linked glycans has been reported, however the functional modification on the terminal Sias was not addressed in detail (38). Here we demonstrate a simple yet versatile encoding technique for the representation of mainly sialyltrisaccharide structures that will help in suitable structural representation of glycans in a machine-readable digital format and hope this will benefit the glyco-community as a whole. The newly developed coding system can also be utilized to calculate the number of possible naturally occurring sialyltrisaccharide structures in mammalian sialomes. Although the complexity and the diversity of the sialome make the calculation of the total number of sialoside structures impossible, we believe that our coding approach will help to address the long-standing question regarding the number of possible glycan structures.

This huge number of sialosides in nature along with their immense role in post-translational modifications and protein-binding make them an important target that needs to be probed for the understanding of structure and function. Plant-based lectins are traditionally used to probe the sialosides, however they are often limited in recognizing diverse and complex sialic acid linkages and modification. In this paper we took advantage of naturally-evolved bacterial toxin B-subunits to develop a universal probe for the terminal sialic acids irrespective of their linkages and modifications. We already addressed the characterization of an AB_5_ toxin B subunit from *Y. pestis* (YpeB) in the preceding paper (accompanying manuscript Khan *et al*.). *Yersinia* genus encompasses a heterogeneous collection of Gram-negative facultative non-sporulating anaerobic bacteria that are divided into different serological groups based on the antibody reactions to different lipopolysaccharides (39). Only three species of *Yersinia, Y. enterocolitica, Y. pestis*, and *Y. pseudotuberculosis*, have been known to infect humans. Enteropathogenic *Y. enterocolitica* is a food-borne pathogen, which is the most prevalent in humans, causing a broad range of gastrointestinal syndromes and may cause septicemia and meningitis (40–42). The extensive sialoglycan microarray study revealed that YpeB binds to various modified Neu5Ac and Neu5Gc-terminated glycans, except when the terminal Sia was modified with an *O*-acetyl group at the C-4 position (preceding manuscript Khan *et al*.). Consequently, the idea of using YpeB as a universal Sia-binding probe had to be dismissed. These shortcomings led us to screen the glycan-binding B subunits of AB_5_ toxins from phylogenetically-related bacteria in search of a broader Sia binding probe. Like *Y. pestis, Y. enterocolitica* also has a very broad range of host compatibility. Besides humans, the bacteria have also been isolated from rodents, domestic animals (e.g., cattle, pigs, dogs, sheep), and other animals like horse and deer (43). A systematic screening of toxin B subunit of *Y. enterocolitica* (YenB) along with the other phylogeny-based bacterial toxin B subunits on the sialoglycan microarray was pursued to discover a universal probe that recognizes almost all types of sialoglycans present in the dynamic mammalian sialome.

## Results and Discussion

### A coding system for individual sialoglycans on the microarray

The importance and coverage of high-throughput glycan microarray technology depend on the number and diversity of glycans present in the specific library. Over the years we developed a library of sialosides for microarrays. A series of natural and non-natural sialoglycans (see SI for full list) were synthesized using chemoenzymatic methods (44). We restricted the sialoside structure primarily in the form of defined trisaccharides, as addressing all of the diverse structures of sialosides in nature is beyond anyone’s scope. Except for a few non-sialosides, the terminal monosaccharides used are Neu5Ac, Neu5Gc, and Kdn along with their natural modifications. The terminal Sias at the non-reducing end are linked with the underlying glycan structures in α2-3, α2-6, and α2-8 fashion. Terminal Sias are linked to the underlying glycan structures that are usually known to occur in nature (precisely in mammals), for example, Type I (Galβ1-3GlcNAcβ), Type II (Galβ1-4GlcNAcβ), Type III or T-antigen (Galβ1-3GalNAcα), Type IV (Galβ1-3GalNAcβ), Type VI (Galβ1-4Glcβ) (45), Lewis antigens, Tn-antigen, sulfated GalNAc, and ganglioside-type structures. Initially, the individual sialoglycans were assigned with a number (Glycan ID) in the order they were synthesized and stored. The assigned glycan IDs listed in the table (SI Table *S1*) are semi-random and do not help in streamlining the sorting and analysis of the microarray data. To address these issues, we developed a new coding system. The purpose of this system was to easily order the glycans in various logical ways during spreadsheet analyses and identifying glycan-binding motifs. Ten-digit machine-readable codes for individual glycans were devised (Table 1). Due to its terminal position, most of the biological interest is in the identity of the Sia. Thus, the coding of the glycans starts from the non-reducing terminal Sias followed by their modifications and linkages. Subsequently, the underlying two monosaccharides (or more) were assigned from the non-reducing end, each with a digit indicating the monosaccharide type, its modifications, and linkage.

**Table 1:**
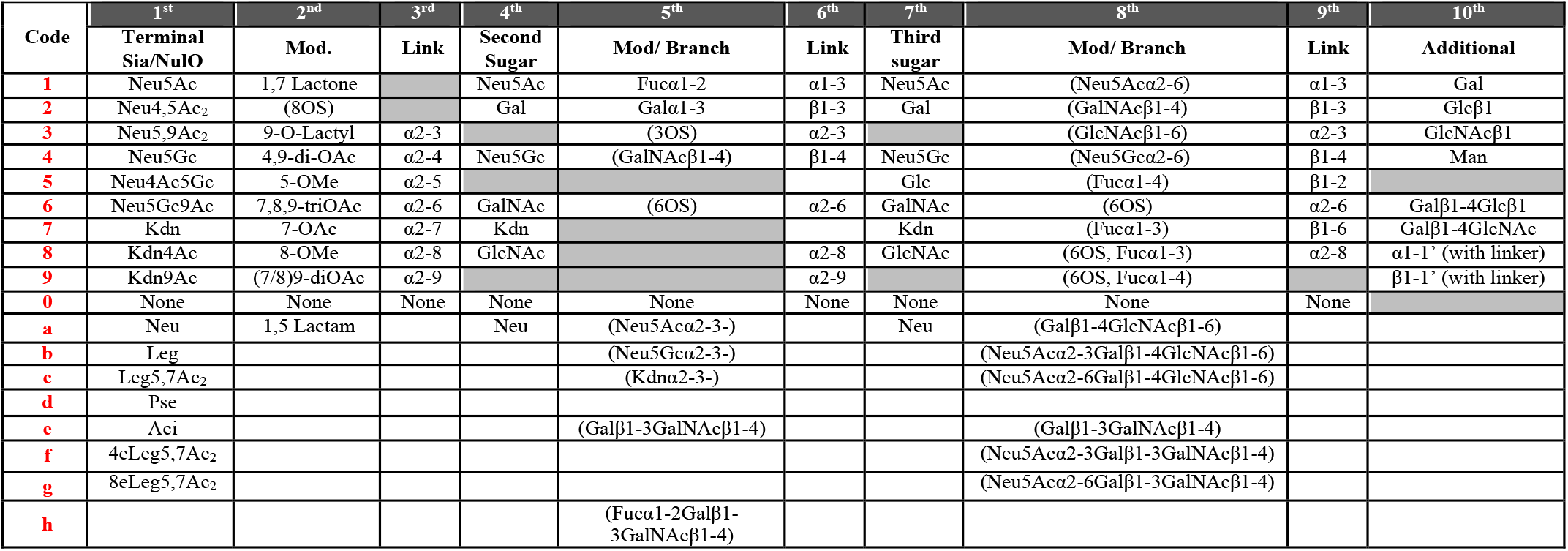
Numerical code for designating naturally occurring trisaccharide sequences carrying terminal sialic acids. In first digit place alphabetical codes are used for less common or bacterial terminal glycan sequences carrying nonulosonic acid units.

The first digit in the 10-digit code represents the terminal Sia identity. Being the most prevalent Sia in mammalian glycomes, Neu5Ac was assigned number *1* followed by its two most important *O*-acetyl modifications at C-4 (number *2*) and C-9 (number *3*) positions. It is noteworthy that Neu5Ac with only C7-*O*-acetyl modification is not stable (46) and hence it was not considered as a primary Sia form. Like Neu5Ac, Neu5Gc and its *O*-acetyl-modified versions were then assigned the number *4* to *6*. The less abundant Kdn and its *O*-acetyl modifications were assigned afterwards to occupy the numbers *7* to *9*. To accommodate the non-sialosides in our library, we decided to assign number *0* for the ‘no sialic acid’ at the non-reducing end of the glycan. We extended the assignment system to accommodate less abundant (or not present in our library yet). An alphabet (small letter) was assigned to each of the neuraminic acid (Neu), legionaminic acid (5,7-di-*N*-acetyl legionaminic acid, Leg5,7Ac_2_), and other nonulosonic acids mostly present in prokaryotes. It is noteworthy that while the existence of ‘naked’ Leg is theoretically possible, all the naturally occurring legionaminic acids reported thus far are *N*-acylated at C-5 and C-7 positions. These sialoglycans were assigned with letters starting from *a* and filling downwards as the library expands. The second digit in the coding system was to identify the modification on the Sia. We assigned numbers *1* through *9* for each type of naturally occurring *O*-acetyl, lactone, *O*-lactyl, 5-OMe, and 8-OMe modifications on the Sia. To the best of our ability, we tried to rationalize the assignment of the number by matching the carbon position of the specific Sia modification. Number *9* was assigned to the (7/8),9-di-*O*-acetyl modification where the position for one of the *O*-acetyl groups is not confirmed due to the *O*-acetyl migration among C-7, C-8, and C-9 positions (46). The 1,5-lactam, a modification generally found in neuraminic acid (Neu) was assigned with alphabet *a*. Next in the column, the third digit represented the various types of natural linkages connecting the C-2 position of Sia to the next underlying monosaccharide unit in alpha stereochemical fashion. To date, no beta linkage was reported to occur at the C-2 position of the Sia (except in the CMP-monosaccharide state). Rationally, the α2-3 connection was assigned to number *3*, α2-6 connection to number *6* and so on. To make our numbering system more self-explanatory, we did not assign number *1* and *2* to any of the linkages, the obvious reason being the absence of α2-1 and α2-2 Sia connections. In total, the first three digits in the coding system were sufficient to properly express most Sias at the non-reducing end. Similarly, the next three digits enumerated the identity, modification/branching, and the linkage of the next underlying monosaccharide. In a poly/oligosialic acid, the Sia is linked to another Sia in α2-8-fashion and for that reason we included Sia in the list of second sugar unit (fourth digit). To keep the numbering system in sync, the Neu5Ac, Neu5Gc, and Kdn in this column were assigned to the same numbers/alphabets as in the first digit place (second column of the Table 1). We assigned leftover numbers to the naturally occurring monosaccharides such as galactose (Gal), *N*-acetylgalactosamine (GalNAc), and *N*-acetylglucosamine (GlcNAc). Apart from the listed ones, the occurrence of any other monosaccharides immediate to the terminal Sia is so far unknown. We did not use the numbers 3, 5, and 9 for the fourth digit place as there are no known linkages of non-terminal sialic acid through C-3, C-5, or C-9 positions (except on some bacterial capsular polysaccharide). Next, the branching at the second sugar unit was represented by the fifth digit. The branching monosaccharides were assigned along with their linkages. The Fucα1-2 (found in Lewis^Y^, Lewis^B^, and human blood group antigens); Galα1-3 (found in B-blood group antigen/alpha-Gal), GalNAcβ1-4 (found in ganglio-series gangliosides), and 3/6-*O*-sulfation were assigned with the numbers that were in sync with the number assignments in other columns. Although we focused on trisaccharides, we took the liberty to accommodate some important natural glycans with longer side chains such as GM1 and Fuc-GM1 structures. The Galβ1-3GalNAcβ1-4 and Fucα1-2Galβ1-3GalNAcβ1-4 branches were assigned with letter *e* and *h* respectively. Next, the sixth digit was assigned to the linkage that connects the reducing end of the second sugar and the non-reducing end of the third sugar unit. The number assignment was in accordance with the previous linkage assignments listed in the fourth column (the third digit). The identity of the third sugar in the linear chain was denoted with the seventh digit. The number assignment of the monosaccharide for the seventh digit place remained the same as in the second sugar unit (fourth digit). The only addition was the glucose (Glc) monosaccharide, which was assigned with number *5*. The branching on the third sugar was designated in the eighth digit place in the coding system. The branched Sias on the third sugar were assigned with the numbers similar to those described for the first and the fourth digits. Again, the naturally occurring glycans and sulfated modifications were assigned with the numbers. In the eighth digit place, a series of di- and tri-saccharide branching (as found in Core 2, GD1) was considered and each was assigned with a letter. The linkage at the reducing end of the third sugar was designated in the ninth digit place. It is important to remember that the linkages had to be between two sugar units. Generally, the third sugar unit in our library was attached with an aliphatic linker at the reducing end and hence the ninth digit was always *0* (none) in our case. We assigned an additional tenth digit place to describe the linker connection or any further monosaccharides that are not present at this moment in our library. The α-linked linker was assigned with number *8* and β-linked linker is assigned with number *9*. Utilizing the coding system, the sialoglycans in our library can be conveniently assigned with a 10-digit code. Fig. 1 demonstrates the representation of sialoglycan in a machine-readable numerical format. See supporting information (Table S1) for the full list of the numerically encoded sialoglycans.

**Figure 1:**
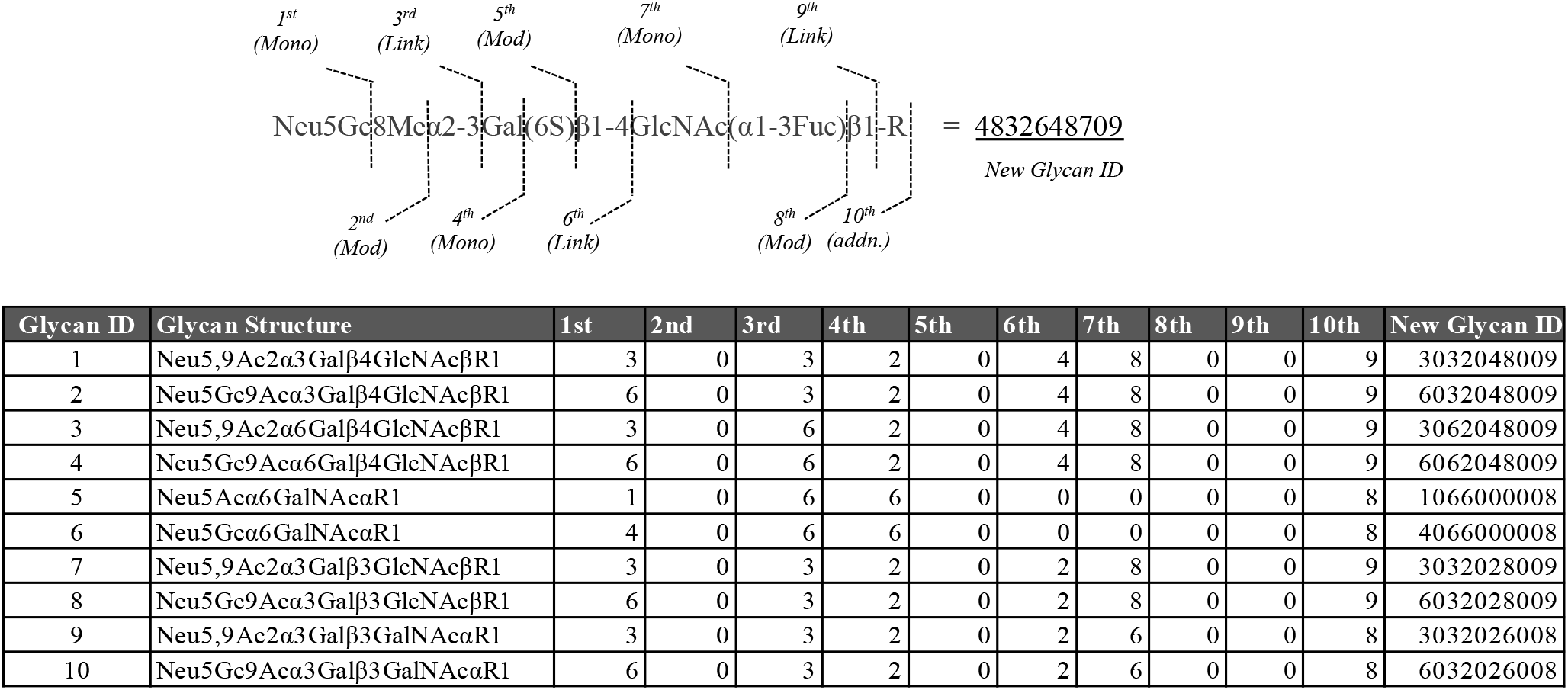
Coding strategy utilized to encode sialyltrisaccharides. Codes associated with the individual glycan identity, modification, and linkage are listed starting from the non-reducing end. Example of 10-digit numerical code representation of sialyltrisaccharides. New glycan id (extreme right column) is generated based on the encoding rules described in Table 1. The left most column describes the semi random old glycan id which is based on the order of which the glycans were synthesized and stored. For complete list see SI Table *S1*.

### Application of the coding system for logical sorting of array results

One of the major reasons behind developing the coding system was to make the microarray data sorting easy and logical. Fig. 2 shows the binding of polyclonal chicken anti-Neu5Gc antibody (**) on the sialoglycan microarray experiment. When the data were plotted against the semi-random glycan ID (Fig. 2A), the interpretation of data became clumsy and lacked a logical explanation. Once we applied the newly developed coding system on the glycans and sorted the data based on the 10-digit code, the chart became very clear and self-explanatory. The 10-digit codes were quickly arranged in an ascending order of the new glycan ID (in Microsoft Excel) to logically group the Neu5Gc-glycans (and its modified versions) together. The data plotted afterwards (as bar diagram) gave a clear visualization of the specificity of the antibody towards the Neu5Gc-epitopes (Fig. 2B).

**Figure 2:**
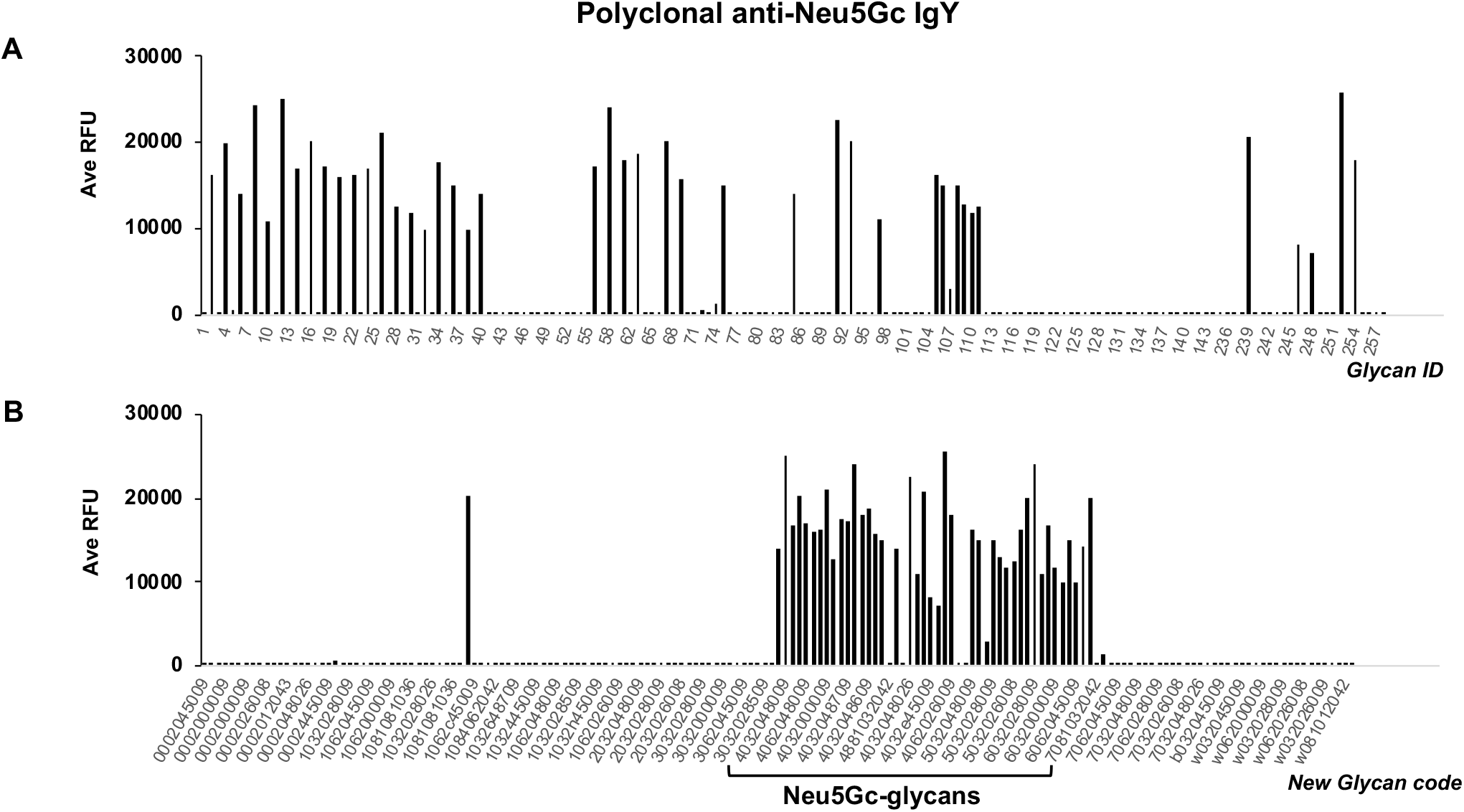
Application of coding system to identify binding patterns. Chicken polyclonal anti-Neu5Gc IgY antibody was tested on sialoglycan microarray platform and average fluorescence unit (Ave RFU) was plotted against a) semi-random glycan id and b) new glycan code arranged according to the terminal Sia identity. The lone strong signal at the left of the plot corresponds to the glycan Neu5Acα6(Neu5Gcα3)Galβ4GlcβR1.

In another example, we demonstrated that the 10-digit code not only can be helpful in arranging the data in ascending or descending order, but the numbers at the individual digit place can also be grouped together to convey a meaningful result. To demonstrate the technique, we tested the biotinylated *Sambucus nigra* lectin (SNA) on the sialoglycan microarray. The binding dataset was organized by grouping the linkages of the terminal Sia unit, i.e., the third digit. From the plot (Fig. 3), it was clearly demonstrated that SNA prefers to bind α2-6-linked Sia epitopes.

**Figure 3:**
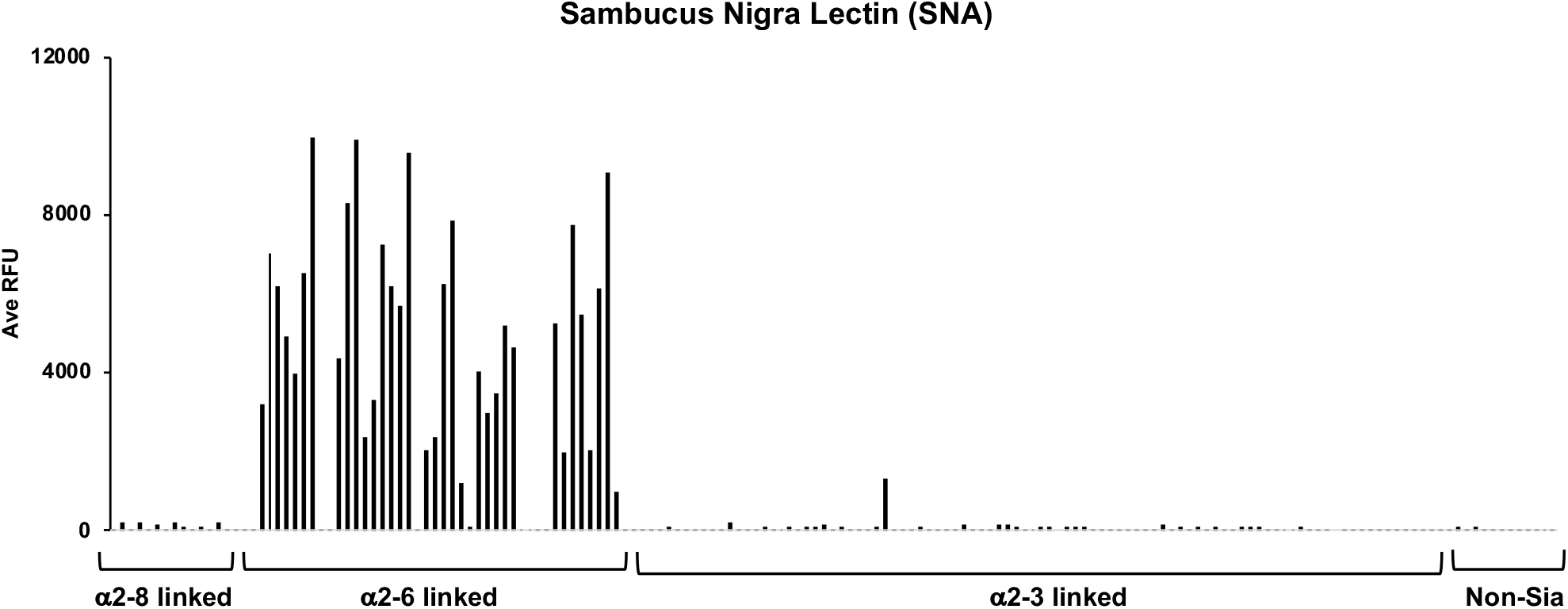
Application of coding system for logical sorting of microarray data. The data obtained from the microarray experiment with biotinylated SNA lectin was organized based on the linkage between the terminal Sia and the underlying glycan structure. The plot conveniently and clearly shows the preference of SNA lectin towards the α2,6-linked Sia.

### Application of the coding system for motif searching of results

Among the versatile applications of the coding system, one important application was to search for the glycan motif responsible for binding to the lectins, proteins, and antibodies. Although our focus was on the terminal Sia, the underlying glycan structure plays a critical role that affects the binding pattern of the glycan-binding biomolecules. It was well-known that *Maackia amurensis* Lectin II (MAL II) recognizes the terminal Sia linked to the underlying glycan structures in α2-3-linkage. The motif finding based on the new coding system also helped us to confirm the dependence of the binding specificity on the underlying glycan structures. We tested biotinylated MAL II on our sialoglycan microarray and a heat-map is plotted (Fig. 4). The top 20 strongest signals (average relative fluorescence unit, Ave RFU) obtained from the MAL II-binding on the microarray were listed along with the numerical code of the sialoglycan epitopes. It became obvious that all the strongly-binding epitopes bore the same number sequence from third to seventh digit places, which eventually indicated that the Sia α2-3 linked to the underlying Galβ3GalNAc (sialyl T-antigen on O-glycan) and Galβ3GlcNAc (type 1) structures serve as the key motif for the MAL II binding. The coded number made the motif recognition much easier compared to finding the motif by reading the structures.

**Figure 4:**
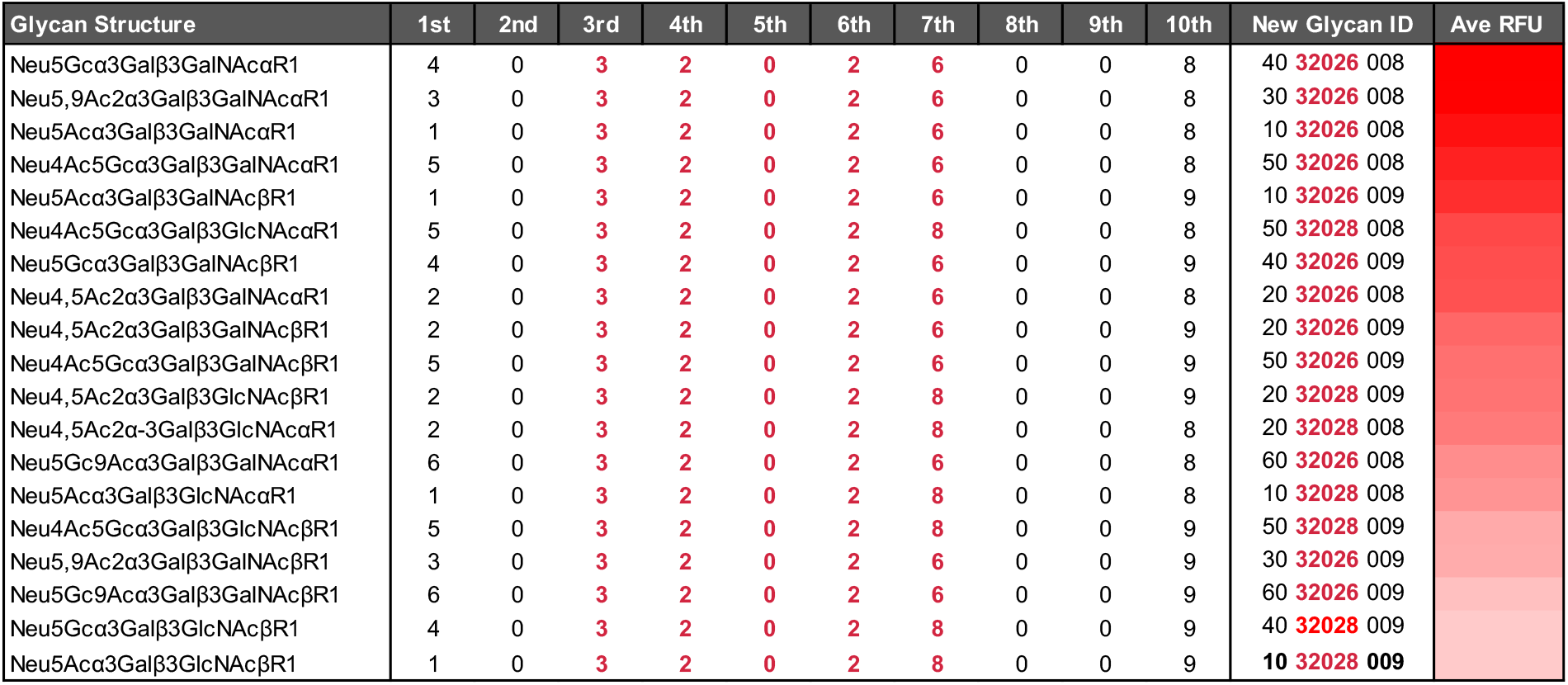
Motif finding utilizing the new sialoglycan code. Top 20 strongest signal obtained from sialoglycan microarray experiment of Biotinylated *Maackia Amurensis* Lectin II (MAL II) was listed along with the 10-digit new glycan id. The digits labelled in red showing a clear motif in glycan structure that serves as the ligand epitope of MAL II. The most prominent motifs determined here are the Siaα3Galβ3GalNAcα (O-glycans) along with the Siaα3Galβ3GlcNAc (type 1) structures.

### Coding-enabled enumeration of sialyltrisaccharide structures plausible and likely to appear in nature

The systematic coding system also helped us calculate the number of possible sialyltrisaccharide structures that could exist in nature. In estimating the space of all, plausible and likely trisaccharides some fundamental concepts were applied. The set of *all* trisaccharides contains every trisaccharide code that can be written down. The set of *plausible* trisaccharides was extracted from that set by reasonably asserting that all sugars must be connected, and no modification or linkage can share a carbon; principles well established in glycobiology. Finally, the *likely* trisaccharides were extracted using a probabilistic approach to generalize glycan motifs found in a set of known trisaccharides. Here, we used co-occurrences observed between each adjacent and non-adjacent glycan element (sugar, linkage, and modification) to determine which two-element motifs were common, observed or never observed. Similar to the studies using motifs to generalize structural feasibility, the joint probabilities were aggregated over each novel trisaccharide to calculate a motif-based likelihood for each plausible trisaccharide. To enumerate all trisaccharide combinations and establish a total number for reference, we used a simple cartesian product; every selection of one element from each of several sets. Here, we enumerated every selection of one of the sugars, linkages, or modifications permissible at each of the 9 positions. The Cartesian product of the trisaccharide code gives 212,889,600 total trisaccharide combinations. To calculate the number of plausible trisaccharides, we selected from the set of all trisaccharides by asserting that trisaccharides must be connected and non-conflicting. Intuitively, no nonreducing sugar may follow a missing linkage (i.e. “None”). Similarly, no linkage or modification may connect to a missing reducing monosaccharide (i.e. “None”). Yet, a missing modification has no impact on the plausibility of the trisaccharide. We also applied constraints specifying the specific carbons each sugar provides for the formation of a glycosidic bond and the carbons required by a linkage or modification. Each monosaccharide provides a discrete number of hydroxyl groups at specific carbons, while each modification and linkage require a free hydroxyl group at specific carbons. In a plausible trisaccharide, each linkage and modification are associated with a sugar providing at least every necessary carbon to support those linkages and modifications. Additionally, the reducing linkage, modification, and nonreducing linkage must not require the same carbons. After removing all unconnected trisaccharides (those with an interceding “None”), all conflicting trisaccharides (those with linkages and modifications requiring the same or unavailable carbons), there are 2,799,962, only 1.32% of the total, plausible trisaccharides remaining. Finally, we ranked the likelihood of trisaccharides by comparing them to a list of 140 known human trisaccharides. Because permissible reaction information is limited for trisaccharides, we used a statistical approach to infer feasible trisaccharide structures from the list of known human structures. We looked at the probability that any two elements in the trisaccharide code would co-occur in the set of 140 known sugars (9 choose 2 = 36 pairs) and used that to generate a log-likelihood scoring. Due to the limited coverage of the combination space, we added a pseudo-count (0.1) to minimize the penalty of pairs unobserved in the known set. Over every plausible trisaccharide, the median log-likelihood was −21.7 and the median number of unobserved pairs per plausible trisaccharide was 6. The least likely trisaccharides have low log-likelihoods (min=-70.9) and high numbers of unobserved pairs (max=34). There were 113,781 trisaccharides with no unobserved pairs and these least unusual trisaccharides had a median log-likelihood of −8.07. There are 471,292 trisaccharides with at most 2 unobserved pairs and 746,861 trisaccharides with at most 3 unobserved pairs. With more unobserved pairs, the likelihood of that trisaccharide naturally occurring decreases.

In counting the total, plausible and likely trisaccharides, we found that these numbers differed by several orders of magnitude. The number of total trisaccharides is useful to orient us to the size of combinatorial possibility but has limited biological meaning as it contains many impossible structures with, for example, 1st and 3rd sugars with no 2nd sugar. The possible trisaccharide structures impose the minimal constraint, the sugar must be connected (no missing links or sugars) and all carbons requested by linkages or modifications must be requested only once and supported by the corresponding sugar. These constraints reduced the total sugar set from over 210 million to just over 2 million. The plausible sugars provide the most forgiving possible constraints and therefore describe an upper-bound suggesting that there are unlikely to be more than 2.8 million trisaccharides. Finally, we imposed a harsh constraint, compliant with a set of known trisaccharides. 113,781 of the plausible trisaccharides were completely compliant with the set of known trisaccharides, containing no code pairs unobserved in the known set, suggesting that there are at least 113,781 sialyltrisaccharides. Though an unobserved pair makes it less likely that a trisaccharide is sterically feasible to assemble, the set of known trisaccharides was not comprehensive. Therefore, a trisaccharide with an unobserved pair may simply be a new type of trisaccharide. The more uncharacterized pairs a trisaccharide has, the less likely it is to contain no impossible code pairs. Therefore, a more forgiving lower-bound could be established as fewer than 2 or 3 unobserved pairs, 471,000 and 747,000 trisaccharides respectively. Within these large steps, trisaccharides can be ranked by log-likelihood to find the most likely sugars within each set. For the >113,000 plausible linear sialyltrisaccharides, we note that a biantennary *N*-glycan with two terminal sialoglycan trisaccharides could have >10^10^ potential combinations and a triantennary *N*-glycan with three terminal sequences, >10^15^ potential combinations at a single *N*-glycosylation site. With this calculation that includes selected sialic acid modifications and linkages, we greatly extended the possibility of the huge number of N-glycans reported previously (47).

### Discovery of phylogenetically related bacterial AB_5_ toxin B subunit from *Yersinia enterocolitica* (YenB) as a universal sialoglycan probe

The huge number of naturally possible sialyltrisaccharide structures, especially when branching is considered, poses an enormous challenge in probing of terminal Sia in a dynamic mammalian sialome. Traditional plant-based lectins are often limited in probing the sialosides due to the diverse and complex linkages and modification of terminal Sia. We addressed this challenge by exploiting the naturally-evolved sialic acid recognizing bacterial proteins which exhibit broad host range. Pathogen-glycan interactions are important both for colonization of host surfaces and recognition and entry of cells by toxins. A broad range of host compatibility (43) and phylogenetic proximity to the *Yersinia pestis* makes *Yersinia enterocolitica* a prime candidate that may express broad Sia recognizing proteins. Earlier studies reported that plasmid-bearing *Y. enterocolitica* adhere to purified small intestinal mucins from rabbits and humans (48, 49). The binding of the virulent bacteria with mucin was inhibited by purified mucin oligosaccharides and thus suggests the involvement of the glycan moiety in the cell specific interaction. Given the phylogenetic proximity of *Y. pestis* and *Y. enterocolitica* (Fig. 5) and the reported binding affinity of *Y. enterocolitica* with mucin carbohydrates, we anticipated that the *Y. enterocolitica* AB_5_ toxin B subunit homologue YenB should recognize diverse Sia epitopes.

**Figure 5:**
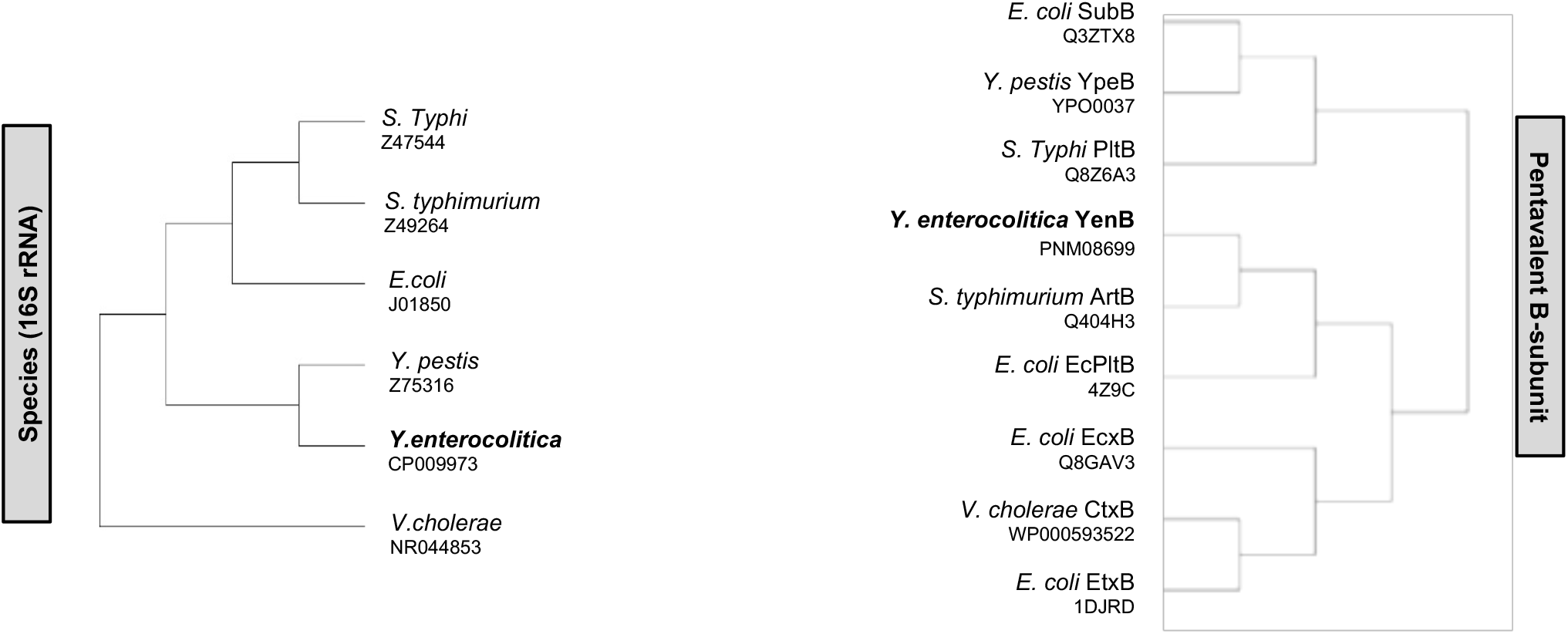
Phylogenetic relationship between species based on 16S rRNA vs pentavalent B subunit of exotoxins. The comparison reveals that the evolution of bacterial AB_5_ toxin B subunit is independent of the species phylogeny. Corresponding accession numbers are indicated below.

YenB (expressed with a C-terminal His6 tag) was purified from recombinant *E. coli* BL21, by Ni^2+^-NTA chromatography. The purified YenB protein was then tested for Sia binding using the sialoglycan microarray platform. Each glycan was printed in quadruplets on a *N*-hydroxysuccinimide coated glass slide and treated with the primary protein of optimized concentration. Average relative fluorescence unit (Ave RFU) was measured after treating with the Cy3-tagged secondary antibody. The sialoglycan coding system was used to rationally sort and plot the data for convenient interpretation. The microarray result (Fig. 6) demonstrated a broader Sia recognition by YenB compared to its phylogenetically-related homologue YpeB from *Y. pestis*. Interestingly, YenB showed a strong binding response against almost all types of Sia in our inventory including the 4-OAc, 5-OMe, and 8-OMe Sia, which YpeB failed to recognize. YenB bound to the Sia irrespective of the linkages (α2-3; α2-6; and α2-8), modifications (4-OAc, 9-OAc, and 8-OMe), and underlying glycan structures. In Fig. 6, the only Neu5Gc-glycan that showed diminished binding was found to be Neu5Gcα3Gal6Sβ4(Fucα3)GlcNAc6Sβ R1. The presence of two sulfate groups probably interferes with the epitope recognition by YenB. The figure legend has been updated indicating this observation. Although YenB recognized the Neu5Ac and Neu5Gc-glycans promiscuously, it did not recognize Kdn-glycans. The lack of binding towards the non-sialosides established that the binding of YenB is not promiscuous towards any glycan epitope.

**Figure 6:**
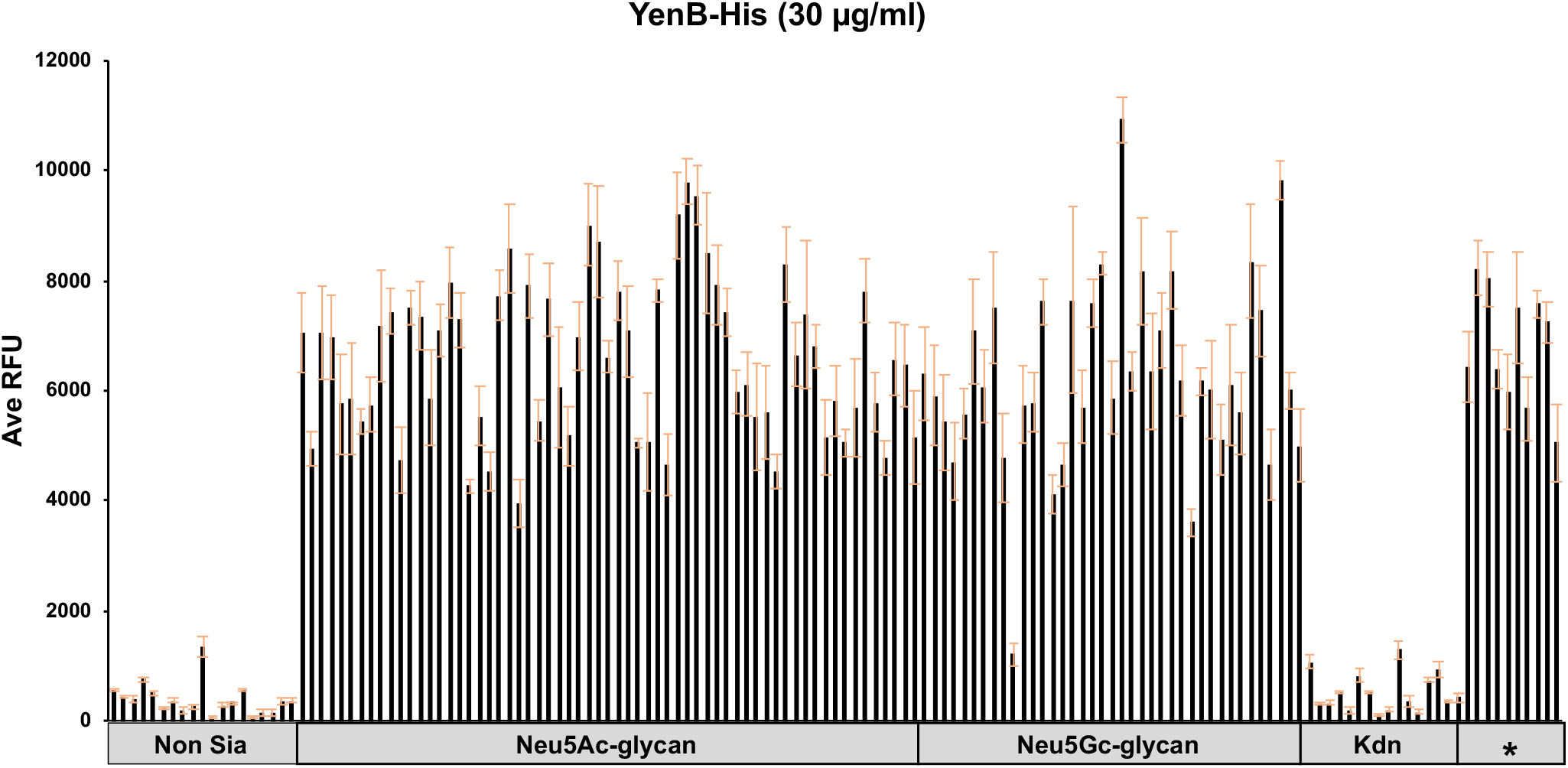
Sialoglycan microarray experiment with B subunit from *Y. enterocolitica*. The chart showed broad range sialic acid recognition by the His-tagged YenB protein (30µg/ml). The glycans on the X-axis is in the order as listed in the Table S1 (supporting information). The non-sialoside and Kdn-glycan epitopes were not recognized by the YenB. Ganglioside type glycan epitopes are designated as asterisk symbol (*) in the chart. Error bar represents the standard deviation for each glycan printed in quadruplicate.

### Structure modeling study of YenB and development of YenB mutant for testing in the microarray

Since the crystal structure of YenB is not yet known, we constructed a structural model of YenB using the crystal structure of SubB (3DWP) as a template. The structure model of YenB was achieved by aligning its primary amino acid sequence with other B pentamers and using crystal structure of SubB as template. The crystal structure of SubB was chosen as template because it shares the highest primary amino acid identity (57%) to YenB. Structural alignment of SubB with YenB (Fig. 7A), and YenB with YpeB (Fig. 7B) showed that YenB adopts the conserved OB fold which is typical of the B subunits from the AB5 toxin family. With the available protein structure of SubB, it is known that serine residues are critical for binding to Neu5Ac, and tyrosine interacts with the extra hydroxyl group of Neu5Gc and are thus critical for binding to sialic acids (50). Upon aligning the sequences of *E. coli* SubB and *Y. enterocolitica* YenB (Fig. 8) we predicted that the mutation of conserved serine residue (S10) might adversely affect the glycan binding preferences. Additionally, the YenB binding site region has isoleucine (I82) as opposed to the conserved tyrosine (Y78) in the SubB (Fig. 9A). Notably, for YenB the residue adjacent to the I82 is a tyrosine (Y81), which is conserved in both species. To ensure a complete disruption of sialoglycan binding by the YenB, we mutated serine to alanine (S10A) and tyrosine to phenylalanine (Y81F) (SI Fig. S1). The recombinant YenB-mutant was expressed, and the protein was purified according to the similar method described for the YenB. The His-tagged YenB-mutant protein was tested on the microarray and no binding with the sialosides or the non-sialosides was observed whatsoever (SI Fig. S2).

**Figure 7:**
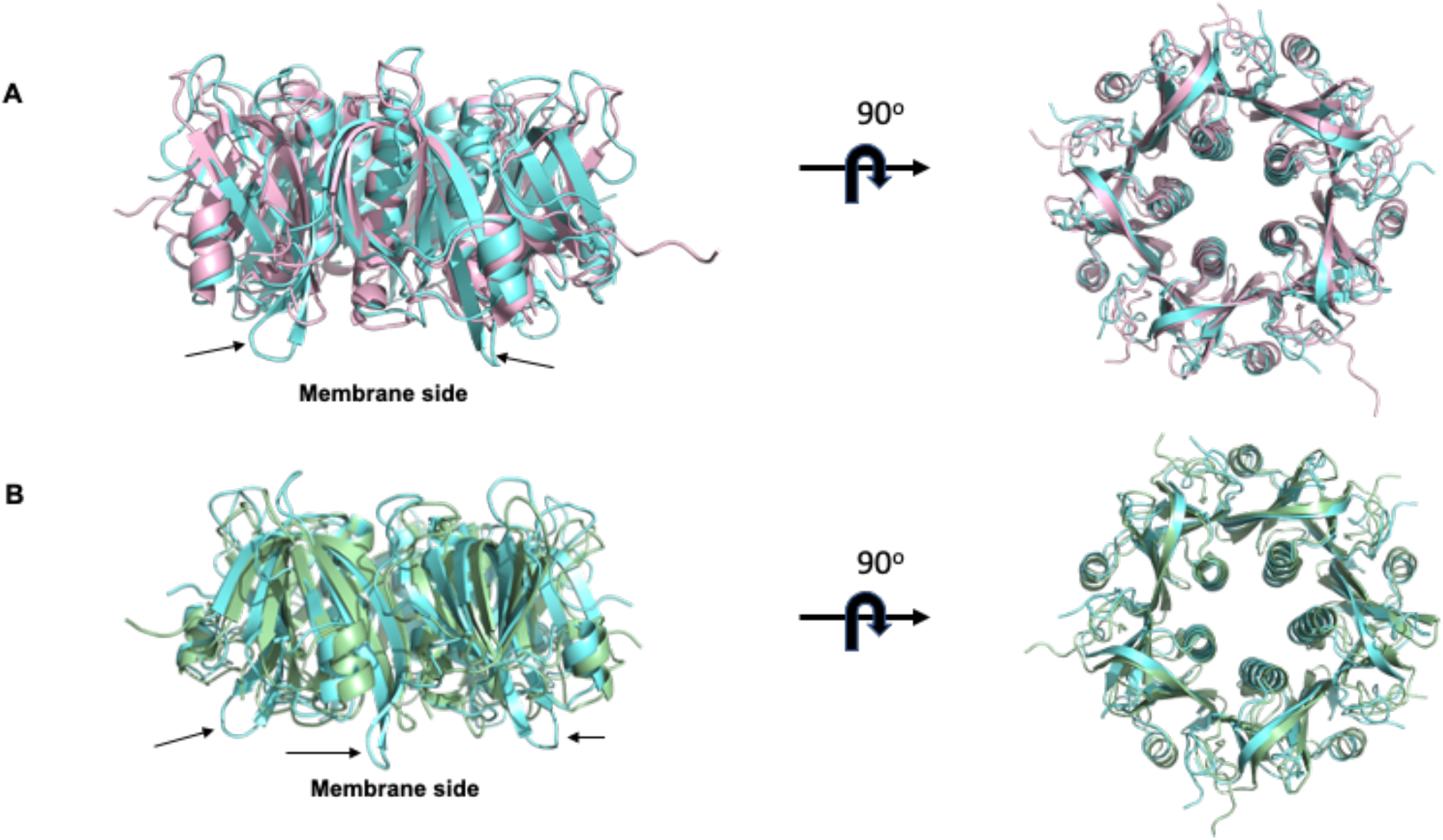
YenB structure modelling and comparison with SubB and YpeB. A) Superposition of the SubB X-ray structure (pale pink) (PDB code:3DWP) with model of YenB (cyan). Overall, very similar to SubB, YenB has longer helix at a distance from the binding site (showed by arrows). B) Superposition of the YpeB (pale green) and YenB (cyan) 3-dimensional model. The structure models are very similar to each other except a few longer helices at a distance from the binding site (showed by arrows).

**Figure 8:**
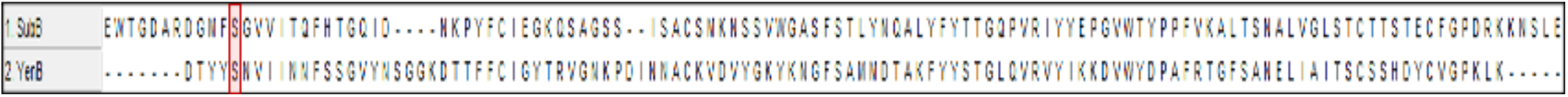
Alignment of the sequence for the B subunits from the *Y. enterocolitica* (YenB) and *E. coli* (SubB). The conserved serine residue is highlighted in red.

**Figure 9:**
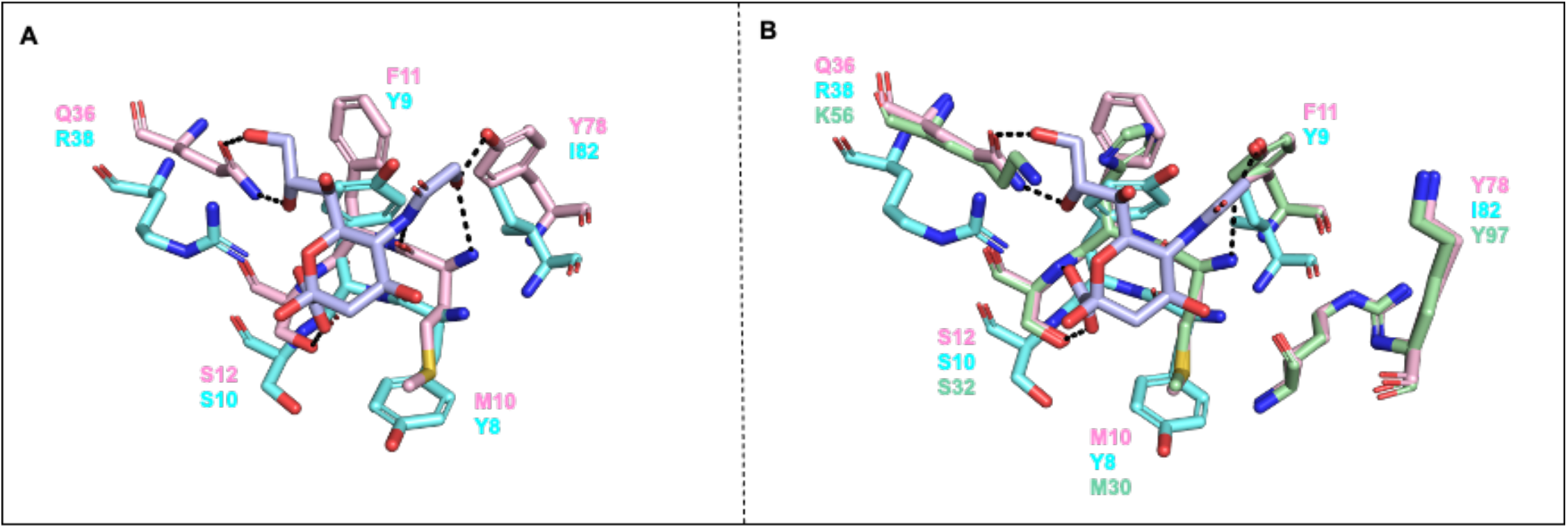
Interaction of sialic acid (Neu5Gc) at the binding pocket of YenB and others. A) Superposition of the Neu5Gc binding site of SubB (pale pink) with the model of YenB (cyan) using the C-alpha coordinates of all residues. Neu5Gc is highlighted in light purple. B) Superposition of the Neu5Gc (light purple) binding site of SubB (pink) with the model of YpeB (pale green) and model of YenB (cyan) using the C-alpha coordinates of all residues. Neu5Gc is highlighted in light purple.

Comparing the Neu5Gc-binding site of SubB with the proposed glycan-binding site within YenB reveals two major differences (Fig. 9A). Firstly, some of the critical SubB residues required for binding are not conserved suggesting that glycan binding is different. Secondly, the glycan-binding site in YenB is located below the position of glycan-binding site in SubB as binding residues in YenB are shifted down. Similar comparison to the YpeB glycan-binding site (Fig. 9B) also revealed a significant departure from the conservation of the amino acid residues at the binding site of YenB. Notably, the YenB has a tyrosine (Y8) residue near the C-4 position of the sialic acid whereas the YpeB and SubB binding pockets have methionine residue (M30 and M10 respectively). This could be a possible reason for the differential 4-OAc sialoglycan binding prefence between YenB versus YpeB and SubB.

### Phylogenetic comparison of YenB with other AB_5_ toxin B subunits (YpeB, SubB, PltB, ArtB, *Ec*PltB, CtxB, and EcxB)

The broad Sia binding propensity of YenB motivated us to explore other phylogenetically-related bacterial AB_5_ toxin B subunits. In the accompanying paper (accompanying manuscript, Khan *et al*.), we briefly described the Sia-binding pattern of the Shiga toxogenic *E. coli* Subtilase cytotoxin B subunit (SubB), which shares 59% identity with YpeB. In addition, Sia-binding preferences of B subunits from closely related pathogenic *Salmonella enterica* serovars, *S*. Typhi (PltB) and *S*. Typhimurium (ArtB) were also tested on the sialoglycan microarray platform. Interestingly, a poor correlation was found between the phylogeny of known B subunit sequences in available genomes and bacterial species phylogeny (based on 16S rRNA). For example, based on B subunit phylogenetic tree inferences, YenB shares its clade with ArtB (54.24% sequence identity) but not with YpeB (Fig. 5).

Other than the aforementioned B subunits, we found three other closely related B subunits (Fig. 5), which might show interesting Sia-binding patterns. Some pathogenic extra-intestinal *E. coli* strains produce a pertussis-like AB_5_ toxin (*Ec*PltAB) (51), related to the toxins from typhoidal and nontyphoidal *Salmonella* serovars. *Ec*PltAB toxin shares 70-80% sequence identity with *S*. Typhi PltAB. The B subunit of the toxin (*Ec*PltB) also shares 69% sequence identity with ArtB from *S*. Typhimurium. A glycan array binding study of *Ec*PltB displayed avidity for a broad range of branched eukaryotic glycans. However, the effect of modifications on the terminal Sia in *Ec*PltB binding was less explored (51).

Cholera is a clinical-epidemiologic syndrome caused by *Vibrio cholerae* that induces severe and at times fatal diarrhea (52), largely attributable to production of cholera toxin (CtxAB), the B subunit of which (CtxB) binds to gangliosides like GM1 (53). It has been reported that the Sia termini on the gangliosides play an important role in target cell recognition (54), although there is also evidence for alternate fucose recognition (55, 56) (which we did not study here). The cholera-like toxins are the largest subgroup of the B_5_ toxin family. A novel cholera-like toxin was identified from a clinical isolate of *E. coli* (EcxAB) derived from diarrheal patients (57). Unlike other cholera-related AB_5_ toxins, the unique mode of pairing of A subunit to the B subunit categorizes the EcxAB toxin in a novel AB_5_ hybrid cholera-like toxin family. Although the A subunits EcxA and CtxA are structurally and functionally very distinct, the B subunits EcxB and CtxB share 63% sequence identity and almost complete structural homology (58). Like CtxB, EcxB was also reported to show strong binding preference towards Neu5Ac-associated ganglioside GM1 and fucosyl-GM1 structures (58). It was therefore of interest to determine whether the binding pattern changes upon substitution of Neu5Ac with Neu5Gc, a naturally less abundant variant which is not present in humans (59).

### Comparison of sialoglycan binding pattern of phylogenically-related B-subunits (YenB, YpeB, SubB, PltB, ArtB, *Ec*PltB, CtxB, and EcxB)

A diverse sialoglycan binding pattern emerged when the phylogenetically-related AB_5_ toxin B subunits were tested on the microarray and the dataset was organized using the coding system. The comparative binding affinity of the phylogenetically-related six B subunits is presented in a standard heatmap format (Fig. 10). A common observation made for all the B subunits (except for EcxB) under consideration was that they did not show any propensity towards the non-sialosides present in our library (SI Fig. *S3A*). Additionally, the B subunits failed to show any affinity towards glycans with a terminal Kdn (a rare mammalian Sia) unless there was an internal Neu5Ac/Gc present in the glycan structure (SI Fig. *S3B*).

**Figure 10:**
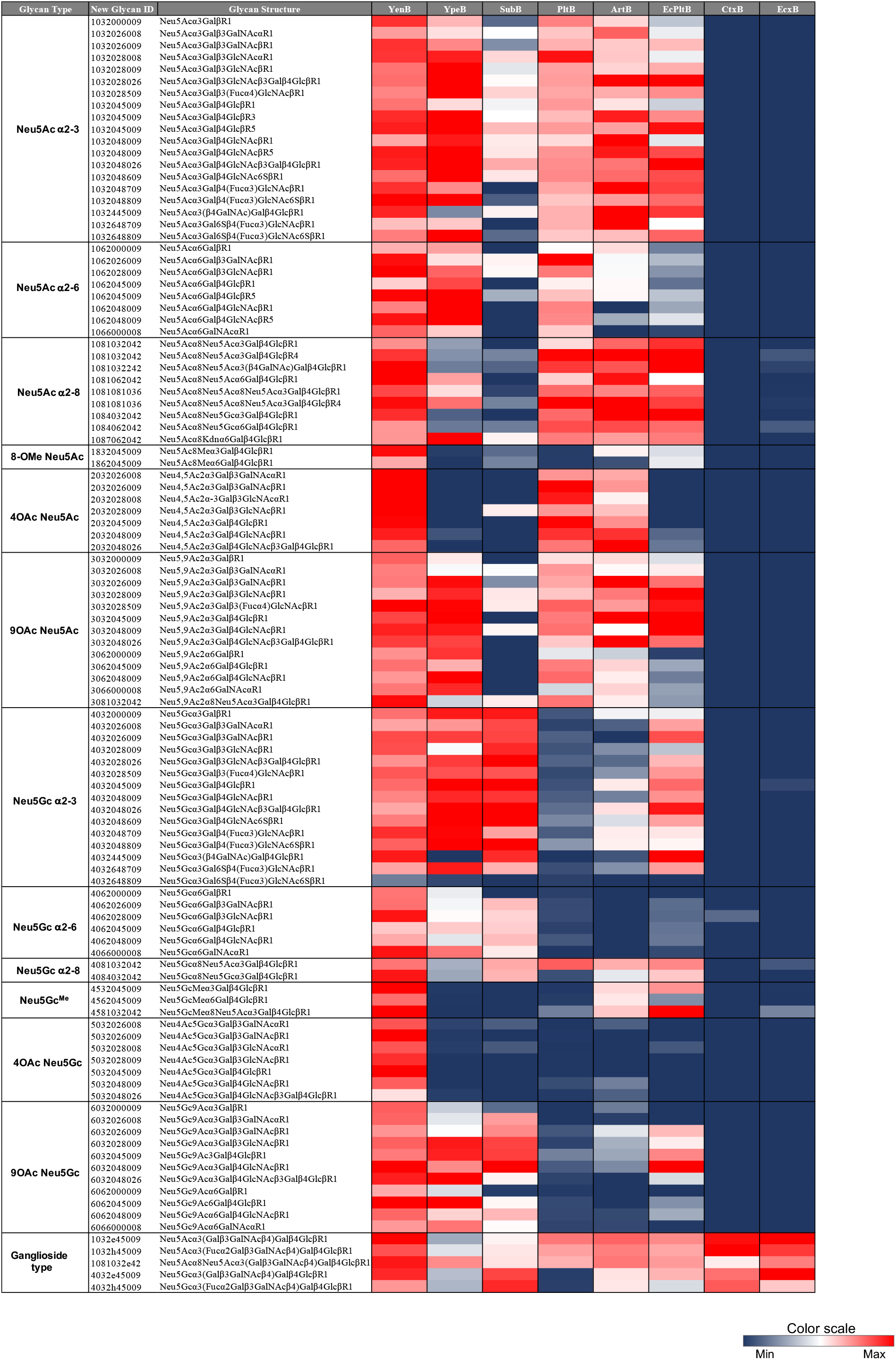
Differential sialoglycan recognition by phylogenetically-related toxin B subunits. Data obtained from sialoglycan microarray experiment showed a diverse Sia binding affinity. The glycan structures were grouped according to the terminal Sia, modifications, and linkages to identify the binding pattern of various phylogenetically-related B subunits. [see supp. Info. section for the Ave RFU numerical data. For a qualitative binding comparison see supp. Table *S2*].

A broad range of Sia-binding by YenB protein has already been demonstrated (Fig. 6). Interestingly, YpeB, a closely related B-subunit protein showed versatile Sia recognition albeit lacking the ability to recognize 4-*O*-acetylated Sia (Fig. 10). The inability of YpeB to bind Neu4,5Ac_2_ and Neu4Ac5Gc may hold the key to answering why some mammals (like horses) are thought to be less susceptible to the plague. Neither of the B subunits could recognize the Kdn-glycans, except when the terminal Kdn is linked to an internal Neu5Ac or Neu5Gc in α2-8 linkage.

A previous report of glycan microarray analysis of *E. coli* SubB showed a stronger binding preference towards the glycans terminating with α2-3-linked Neu5Gc compared to its Neu5Ac congener (50). Testing of SubB in the expanded sialoglycan microarray revealed a similar binding pattern with a preferential α2-3-linked Neu5Gc binding, irrespective of the underlying glycan structures. A weaker albeit significant binding was observed towards α2-6 and α2-8-linked Neu5Gc. The modification on Neu5Gc showed an immense effect in SubB binding. A moderately strong binding was observed towards the 9-OAc Neu5Gc-glycans, however the 5-OMe and 4-OAc modified Neu5Gc completely eliminated the affinity towards SubB. The loss of ligandability of Neu5Gc upon 4-O-acetylation reaffirms the structural similarity of SubB to YpeB, which shares high sequence homology. Notably, Kdn associated glycans did not exhibit any binding to SubB.

Human specific typhoid toxin B subunit (PltB) binds exclusively with Neu5Ac-glycans (60). A detailed analysis of the binding preferences showed that the binding intensity decreases moderately when the terminal Neu5,9Ac_2_ is linked to the underlying glycan in α2-6 linkage. Interestingly, the binding was completely diminished when the terminal Neu5Ac was protected with an 8-OMe group.

An interesting observation was made when ArtB was tested on the microarray. Relative to PltB, ArtB showed a broader Sia preference with strong propensity towards α2-3 and α2-8-linked Neu5Ac- and acetylated Neu5Ac-glycans. However, terminal Neu5Ac with α2-6-linkage exhibited 3 to 5-fold less binding intensity towards ArtB. Sialosides with terminal Neu5Gc showed a mild detrimental effect in binding with ArtB, which is unlike PltB where the Neu5Gc-glycan completely blocks Sia recognition.

*Ec*PltB has been reported to preferably bind with Neu5Ac-glycans when tested at a higher concentration (100µg/ml) but partially loses its selectivity at low concentration (1 µg/ml) (51). We tested the protein at 30 µg/ml concentration and a mild preference towards Neu5Ac over Neu5Gc-glycans was observed. However, the Sia linkage plays an important role in binding affinity. *Ec*PltB showed a clear preference towards α2-3 and α2-8-linked Sia and failed to bind with α2-6-linked Sia, a distinct trait that matches that seen in ArtB. Additionally, Sia recognition by *Ec*PltB was completely blocked by 4-OAc modified Neu5Ac/Gc; a similar observation was made for YpeB and SubB.

The glycan recognition by CtxB and EcxB was very different when compared to other toxin B subunits under consideration. It is known that CtxB preferentially binds to naturally occurring GM1, where the Sia presented is Neu5Ac (53). When tested on the sialoglycan microarray we found that CtxB distinctly recognized both Neu5Ac- and Neu5Gc-associated GM1, as well as fucosylated-GM1. As expected, the binding intensity greatly diminished with GD1 as a ligand. No other sialosides and non-sialosides showed any affinity towards CtxB.

Like CtxB, EcxB also displayed a strong binding preference towards Neu5Ac- and Neu5Gc-GM1. It also recognized the fucosylated-GM1 glycan, albeit with relatively weaker affinity. Like in CtxB, GD1 was also found to be a moderately weak receptor for EcxB. The most striking differences in glycan binding between CtxB and EcxB was the recognition of non-sialosides. Notably, EcxB but not CtxB recognized the non-sialosides present in our library, which represented the terminal fragments of glycosphingolipids of either lacto, neo-lacto, or ganglio-series (Fig. S3). Fig. 10 heatmap shows a comparison of Sia-binding preferences.

## Conclusions and Perspectives

Sialoglycan microarray analysis presented an immense advantage in identifying novel interactions between sialoglycans and other biomolecules. We have established a chemoenzymatically synthesized defined sialyltrisaccharide library for the microarray technology platform. The challenges in sorting and finding motifs from vast microarray data were addressed by developing a novel 10-digit alpha-numerical coding system. The assignment of the code to the individual monosaccharide, their modification, and linkage made the analysis and visualization of data much easier and streamlined. This machine-readable code can be used to find binding target motifs. Using this coding system, we attempted to calculate the number of plausible sialyltrisaccharide structures in nature. Our calculation revealed a remarkable number of 10^5^ possible linear structures of sialyltrisaccharides and hence gave an idea of the huge structural diversity in whole sialomes where the structures of post-translational glycan modifications are not always linear.

We evaluated the binding preferences of phylogenetically-related bacterial AB_5_ toxin B subunits towards the sialoglycans using the sialoglycan microarray platform. We identified that the B subunit from *Y. enterocolitica* could recognize all Neu5Ac and Neu5Gc glycans present in our library irrespective of their modifications and linkages. This gives us an opportunity to use this toxin B subunit as a universal sialoglycan probe. We also re-evaluated the sialoside binding affinity of other phylogenetically-related B subunits. They presented a diverse array (SI Table *S2*) of Sia-binding preferences, which can be exploited to develop a set of sialoglycan binding probes. We realize that it is impossible to have a microarray library that includes every possible naturally occurring sialoside. Moreover, the complexity of cell surface sialomes is beyond the scope of the defined sialoglycan library. To probe the immense diversity of sialosides in biological models, we propose the development of Sialoglycan Recognition Probes (SGRP) (See Srivastava et al. accompanying manuscript).

## Experimental Procedures

### Enumeration and estimation of trisaccharides

In estimating the space of all plausible and likely trisaccharides some fundamental concepts were applied. The set of *all* trisaccharides contains every trisaccharide code that can be written down. The set of *plausible* trisaccharides was extracted from that set by reasonably asserting that all sugars must be connected, and no modification or linkage can share a carbon; principles well established in glycobiology. Finally, the *<u>likely</u>* trisaccharides were extracted using a probabilistic approach to generalize glycan motifs found in a set of known trisaccharides.

In a plausible trisaccharide, each linkage and modification are associated with a sugar providing at least every necessary carbon to support those linkages and modifications. Additionally, the reducing linkage, modification, and nonreducing linkage must not require the same carbons. Intuitively, no nonreducing sugar may follow a missing linkage (i.e. “None”). Similarly, no linkage or modification may connect to a missing reducing monosaccharide (i.e. “None”). Yet, a missing modification has no impact on the plausibility of the trisaccharide. We also applied constraints specifying the specific carbons each sugar provides for the formation of a glycosidic bond and the carbons required by a linkage or modification. Each monosaccharide provides a discrete number of hydroxyl groups at specific carbons, while each modification and linkage require a free hydroxyl group at specific carbons.

To calculate likely trisaccharides, we looked at the probability that any two elements in the trisaccharide code would co-occur in the set of 140 known sugars (9 choose 2 = 36 pairs) and used that to generate a log-likelihood scoring. Due to the limited coverage of the combination space, we added a pseudo-count (0.1) to minimize the penalty of pairs unobserved in the known set.

### Expression and purification of *Yersenia enterocolitica* B-subunit (YenB)

The method described in accompanying paper (accompanying manuscript Khan *et al*.) was followed to express and purify YenB protein with a C-terminal His_6_ tag. The codon-optimized open reading frame encoding YenB was chemically synthesized, flanked by an *Eco*RI site and a ribosome binding site 5’ to the initiating ATG codon, and with a sequence encoding a His_6_ tag fused to the C-terminus of the ORF and a 3’ *Hin*dIII site, cloned into the respective sites of pUC57 (GenScript; Piscataway, NJ). The fragments were then excised and cloned into the *Eco*RI and *Hin*dIII sites of the expression vector pBAD18 (61) such that the ORF is under the control of the vector *ara* promoter. This construct was then transformed into *E. coli* BL21.

For protein purification, the recombinant bacterium was grown in 500 ml LB Broth supplemented with 50 µg/ml ampicillin at 37°C to late logarithmic phase, diluted in half with fresh medium supplemented with 0.2% arabinose to induce *yenB* expression, and then incubated overnight at 26°C. Cells were harvested by centrifugation, resuspended in 20 ml loading buffer (50 mM sodium phosphate, 300 mM NaCl, 20 mM imidazole, pH 8.0), and lysed in a French pressure cell. Cell debris was removed by centrifugation at 20,000 × *g* for 30 min at 4°C. The supernatant was then loaded onto a 2 ml column of Ni-nitrilotriacetic acid (NTA) resin, which had been preequilibrated with 20 ml loading buffer. The column was then washed with 40 ml loading buffer, and bound proteins were eluted with a 30-ml gradient of 0 to 500 mM imidazole in loading buffer; 3 ml fractions were collected and analyzed by SDS-PAGE, followed by staining with Coomassie blue or Western blotting with monoclonal anti-His_6_. Fractions containing >95% pure YenB were pooled, concentrated and stored in PBS/50% glycerol at −15°C.

### Modeling study of YenB with sialosides

The YenB and other amino acid sequences were aligned using the program EXPRESSO (62), and homology modeling was performed employing MODELLER software (63), using the crystal structure of SubB (3DWP) as a template.

### Sialoglycan microarray

The sialoglycan microarray method was adapted and modified from the literature reported earlier (36, 64). Defined sialosides with amine linker were chemoenzymatically synthesized and then quantitated utilizing an improved DMB-HPLC method (46). 100 µM of sialoglycan solution (in 300mM Na-phosphate buffer, pH 8.4) was printed in quadruplets on NHS-functionalized glass slides (PolyAn 3D-NHS; catalog# PO-10400401) using an ArrayIt SpotBot^®^ Extreme instrument. The slides were blocked (0.05M ethanolamine solution in 0.1M Tris-HCl, pH 9.0), washed with warm Milli-Q water and dried.

The slides were then fitted in a multi-well microarray hybridization cassette (ArrayIt, CA) to divide into 8 wells and rehydrated with 400 µl of Ovalbumin (1% w/v, PBS) for one hour in a humid chamber with gentle shaking. After that the blocking solution was discarded and a 400 µl of solution of the toxin B-subunit (30 µg/ml) in the same blocking buffer was added to the individual well. The slides were incubated for 2h at room temperature with gentle shaking and the slides were washed with PBS-Tween (0.1% v/v). The wells were then treated with Cy3-conjugated anti-His (Rockland Antibodies & Assays; Cat# 200-304-382) secondary antibody at 1:500 dilution in PBS. Followed by gentle shaking for 1 hour in the dark and humid chamber. The slides were then washed, dried, and scanned with a Genepix 4000B scanner (Molecular Devices Corp., Union City, CA) at wavelength 532 nm. Data analysis was performed using the Genepix Pro 7.3 software (Molecular Devices Corp., Union City, CA). The raw data analysis and sorting using the numerical codes were performed on Microsoft Excel.

## Data availability

All data are contained within the main text and supporting information.

## Acknowledgments

This work was supported by NIH Grant R01GM32373 (to A.V.), the Novo Nordisk Foundation (NNF10CC1016517, NNF20SA0066621), and NIGMS R35GM119850 (to N.E.L).

